# Modeling dry eye with an air-liquid interface in corneal epithelium-on-a-chip

**DOI:** 10.1101/2023.10.08.561437

**Authors:** Rodi Kado Abdalkader, Romanas Chaleckis, Takuya Fujita, Ken-ichiro Kamei

## Abstract

Dry eye syndrome (DES) is a complex ocular condition characterized by an unstable tear film and inadequate tear production, leading to tissue damage. Despite its common occurrence, there is currently no comprehensive *in vitro* model that accurately reproduce the cellular characteristics of DES. Here we modified a corneal epithelium-on-a-chip (CEpOC) model to recapitulate DES by subjecting HCE-T human corneal epithelial cells to an air-liquid (AL) interface stimulus. We then assessed the effects of AL stimulation both in the presence and absence of diclofenac (DCF). Transcriptomic analysis revealed distinct gene expression changes in response to AL and AL_DCF, affecting pathways related to development, epithelial structure, inflammation, and extracellular matrix remodeling. Both treatments upregulated *PIEZO2*, linked to corneal damage signaling, while downregulating *OCLN*, involved in cell-cell junctions. They increased the expression of inflammatory genes (e.g., *IL6*) and reduced mucin production genes (e.g., *MUC16*), reflecting dry eye characteristics. *TGFB1*, crucial for corneal wound healing, was slightly downregulated in AL_DCF, potentially affecting wound healing processes rather than reducing inflammation by DCF. Metabolomic analysis showed increased secretion of metabolites associated with cell damage and inflammation (e.g., methyl-2-oxovaleric acid, 3-methyl-2-oxobutanoic acid, lauroyl-carnitine) in response to AL and even more with AL_DCF, indicating a shift in cellular metabolism. This study showcases the utilization of AL stimulus within the CEpOC as a comprehensive approach to faithfully reproduce the cellular characteristics of DES.

## Introduction

Dry eye syndrome (DES) is a prevalent and serious ocular condition that has a significant impact on the quality of life of affected patients. The condition is characterized by a range of symptoms, including ocular discomfort, inflammation, reduced visual acuity, and tear film instability, and its estimated prevalence varies between 20-30 % depending on the population and diagnostic criteria used^12^. There are two main subtypes of DES, evaporative dry eye caused by decreased production of the lipid layer of the tear film and aqueous deficient dry eye caused by reduced production of the aqueous layer of the tear film^34^. The underlying causes of each subtype are complex and multifactorial, and include biological, environmental, and physiological factors^5^.

Despite its prevalence and impact, the study of DES has been hindered by a lack of suitable *in vitro* models that can accurately capture the biological and biomechanical aspects of the disease. Conventional *in vitro* models such as cell culture inserts known as transwells, lack the capability to simulate the complex microenvironment of the ocular surface, underscoring the full understanding of disease progression and underlying mechanisms^6,7^. Additionally, *in vivo* animal models are limited in their ability to provide insights into the molecular mechanisms of disease development in human^8^.

To overcome these limitations, organ-on-a-chip (OoC) technology offers an appealing solution for the development of *in vitro* models ^9^. OoC technology provides several advantages, including the ability to control cellular architecture and the capacity to recapitulate external stimuli such as air flow or liquid shear stress^9^, which are essential factors in DES development.

We previously reported the development of a microfluidic device that was used as an OoC for growing corneal epithelial cells under the application of cycles of liquid stimuli for mimicking the eye blinking process^10^. Our previous studies have demonstrated the effectiveness of this technology in recapitulating tear movement during eye blinking and providing temporal analysis of extracellular metabolites through a wide range of corneal transporters^11^. Moreover, OoC of the human cornea with a blinking lid was developed by a different group to model DES^12^. This was achieved by reducing the blinking rates, to allow a tear-like evaporation process; however, the specific mechanism concerning the changes in cell phenotypes under these conditions has not been fully addressed.

Considering that the pathology of corneal damage in evaporative DES is influenced by multiple factors, including tear film disturbances due to eye blinking and exposure to air. Therefore, creating a model that combines both air and liquid interfaces can potentially provide a more realistic representation of evaporative DES in the cornea. However, the extent to which the application of air-liquid stimuli in a microfluidic device can induce a DES-like phenotype in corneal epithelial cells remains unclear. Further investigation is necessary to address this question comprehensively and enhance our understanding of the effects of these stimuli on homeostasis of the corneal epithelium.

Here, we developed an air-liquid interface (AL) system within the CEpOC to mimic the DES-like cellular phenotype. Subsequently, we applied AL in combination with Diclofenac (DCF), a well-known non-steroidal anti-inflammatory drug with a controversial efficacy and safety profile in DES treatment. To gain insights into the underlying molecular mechanism, we conducted a comprehensive dual-omics analysis, including RNA-sequencing (RNA-seq) of corneal epithelial cells and non-targeted LCMS-based analysis of extracellular metabolites collected spatiotemporally from microfluidic devices. Our research provides a better understanding of the cellular responses and metabolic changes associated with AL and with DCF treatment in the context of DES.

## Results

### Creating AL interface in microfluidic devices via CEpOC modification

We have made modifications to CEpOC device in order to generate AL stimuli. This entailed the creation of an AL interface within the microfluidic device. Our approach involved introducing a new component: an open reservoir with a diameter of 4 mm, positioned at the outlet area **(Figure 1. A, B)**. For the generation of air source, we integrated a syringe pump with an air compressor tube. This tube branches into eight separate channels which are subsequently connected to each inlet of the microfluidic device. By employing a two-way flow system involving withdrawal and infusion, we were able to apply two cycles of stimuli. During the first cycle, air is introduced through the outlet, where it effectively blends with the liquid present within the cell culturing channels. This establishes the initial conditions for generating the AL interface. In the subsequent cycle, the air is expelled, allowing for the replacement of the liquid. A visual representation of this process can be found in the supplementary figure **(Supplementary figure 1. A, B, C).**

**Figure 1.**
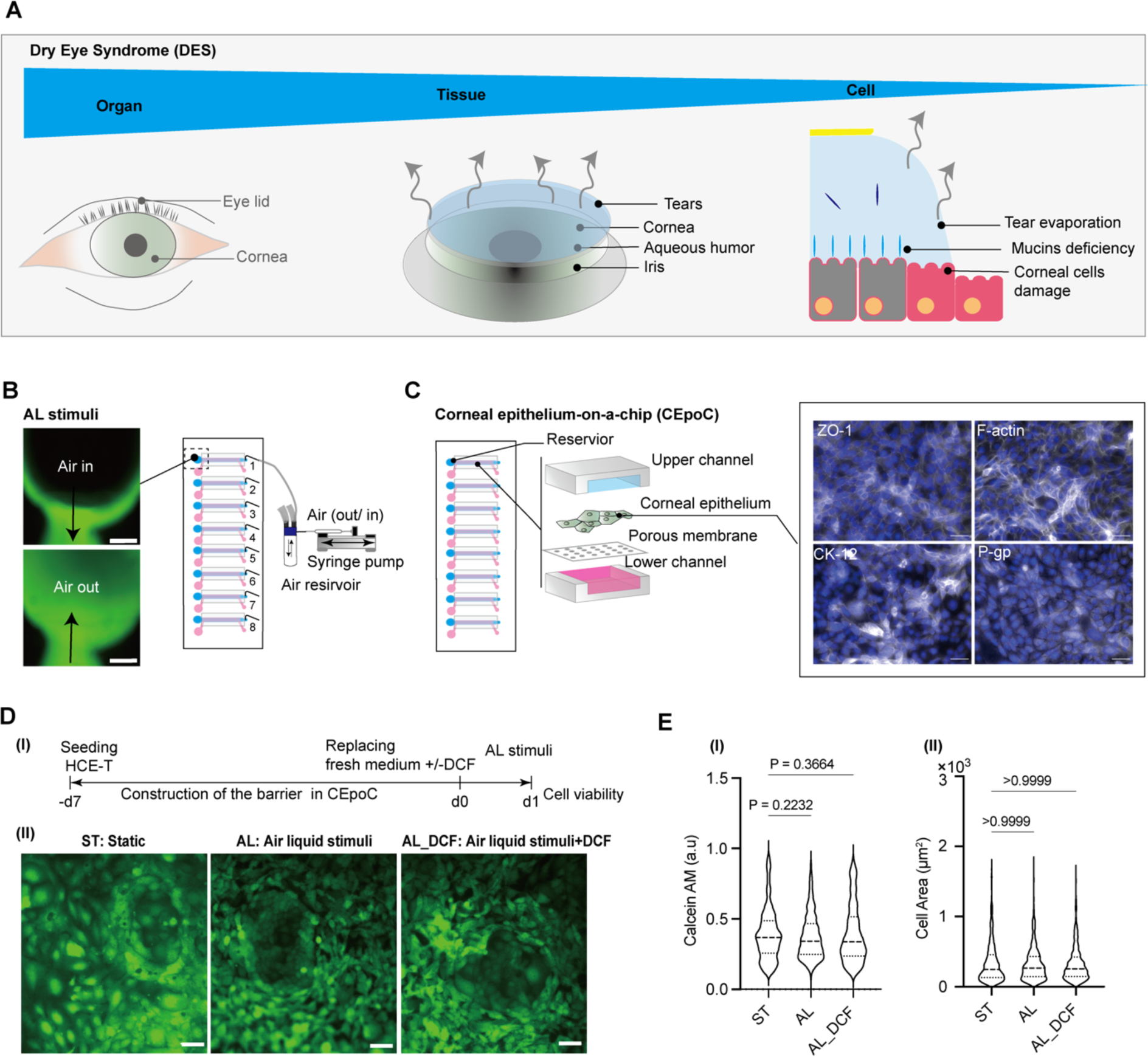
The conceptual design and characteristics of the modified CEpOC. (A) Schematic illustration showcases the evaporative dry eye. (B) The generation of AL stimulus in the modified CEpOC. Green color indicates the fluorescent signal of sodium fluorescein solution in the reservoir area. Scale bar, 500 μm. (C) HCE-T culture at the microfluidic device forming a barrier that express the tight junction protein ZO-1, F-actin, CK12, and P-gp. White color indicates the fluorescent signal of the targeted protein. Blue color indicates the nucleus satin DAPI. Scale bar, 50 μm. (D) Cell viability in the microfluidic devices under ST and AL conditions with and without DCF; I) The protocol that showcase of HCE-T culturing, barrier formation and the AL stimuli; II) Microscopic images of Calcein-AM under ST and AL conditions with and without DCF. (E) Microscopic signal-cell analysis of Calcein-AM intensity (I) and cell area (II). The analysis was performed on images of three independent samples; 1000 cells were randomly selected and analyzed for each sample. Green, Calcein-AM. Scale bar, 50 μm. Data are represented in the violin plot in which the median of each group is indicated with a scattered line (25^th^ 75^th^ interquartile range). The p-values were determined by the Tukey’s multiple comparison test.

### Constructing the corneal epithelial barrier in the modified CEpOC

To establish the corneal epithelial barrier, we followed our previously established protocol employing the human corneal epithelial cell line (HCE-T). The cells were cultivated for seven days, during which the formation of the epithelial barrier was established.

Confirmation of the epithelial barrier’s presence was accomplished through immunofluorescence staining of the tight junction protein known as zonula occludens-1 (ZO-1), in conjunction with the visualization of actin fibers. This indicated the formation of the barrier. Additionally, the cells exhibited other important characteristics, including the expression of the maturation marker cytokeratin 12 (CK12), along with the presence of the apical marker, P-glycoprotein 1 (P-gp) (**Figure 1. C)**.

### AL stimulus exhibits no significant impact on cell viability

To assess the effects of AL alone and in combination with DCF, we subjected the devices to AL stimulus cycles for a duration of 24 h **(Figure 1. D(I))**. Subsequently, we performed cell viability evaluations using Calcein AM staining, comparing the results with devices that were maintained under static control conditions with only the cell culturing medium. Our findings revealed that there was no significant decrease observed in both cell viability presented in Calcein AM intensity and cell area when exposed to either AL or in the combination with DCF (AL_DCF), in comparison to control cells that were kept under static conditions **(Figure 1. D(II)) (Figure 1. E)**.

### AL elicit transcriptomic changes related to corneal function, developmental process, and inflammatory pathways

To evaluate the impact of AL stimulus on corneal epithelial cells, we conducted RNA-seq analysis after subjecting the cells to either AL alone or in combination with DCF for 24 h (**Figure 2. A (I)**). To investigate the transcriptome changes that occur during the activation of biological pathways in response to different stimuli, we first performed cluster analysis of gene expression data. The results distinctly revealed alterations in gene expression patterns in response to AL stimuli alone or in combination with DCF, as compared to the control groups (ST and ST_DCF) (**Figure 2. A (II)**).

**Figure 2.**
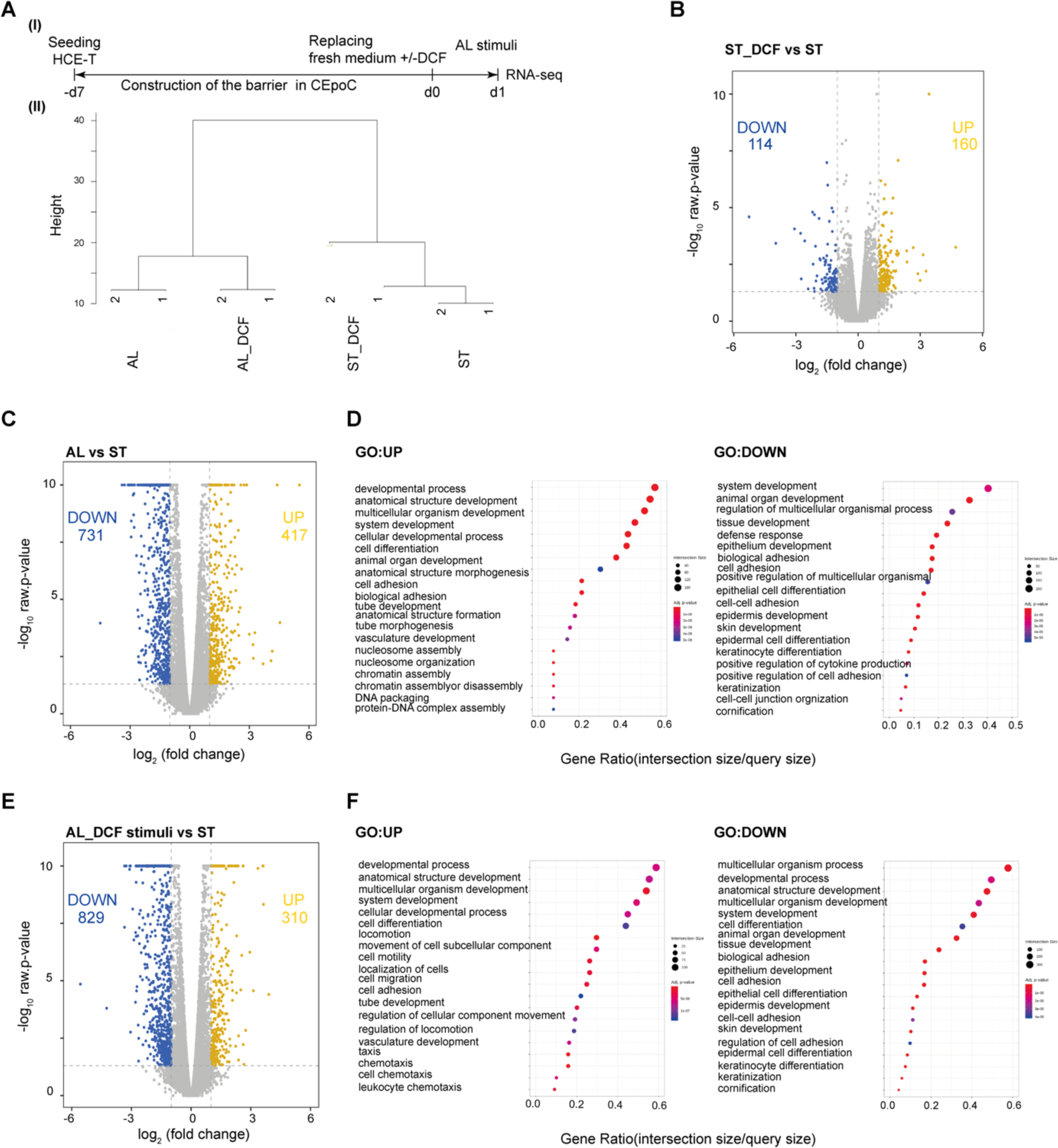
RNA-seq analysis of cells under AL stimulus. (A) I) The protocol that showcase the timeline of HCE-T culturing, barrier formation, and samples collection for RNA-seq; II) Dendrogram indicating the hierarchical clustering analysis by showing the expression similarities among ST, ST_DCF, AL, and AL_DCF (Distance metric = Euclidean distance, Linkage method = Complete linkage). (B) The volcano plot representing the differentially expressed genes (DEGs) of ST_DCF vs ST. (C) The volcano plot representing the DEGs of AL vs ST. (D) GO pathways of DEGs in AL vs ST. (E) The volcano plot representing the DEGs of AL_DCF vs ST. (F) GO pathways of DEGs in AL_DCF vs ST. Genes with 2-fold change and p-value <0.05 was applied. Blue: downregulated genes, yellow: upregulated genes.

To identify the differentially expressed genes (DEGs), we conducted a DEG analysis between the control groups (ST_DCF versus ST) and the experimental groups (AL versus ST and AL_DCF versus ST). In the DEG analysis of ST_DCF versus ST, we identified 274 DEGs, with 160 upregulated genes and 114 downregulated genes (**Figure 2. B)**. The significantly upregulated genes were only annotated with the GO pathway of vitamin D 24-hydroxylase activity (GO:0070576) (**Supplementary figure 2**). In contrast, the DEG analysis of AL versus ST showed 1148 DEGs, with 417 upregulated genes and 731 downregulated genes (**Figure 2. C)**. The significantly upregulated genes were annotated with GO pathways related to developmental processes (GO:0032502), anatomical structure development (GO:0048856), cell differentiation (GO:0030154), and cell adhesion (GO:0007155). The downregulated genes were annotated with various GO pathways related to tissue development (GO:0009888), defense response (GO:0009653), epithelial development (GO:0060429), regulation of cytokine production (GO:0001819), keratinization (GO:0031424), and cornification (GO:0070268) (**Figure 2. D)** (**Supplementary figure 3**).

In the DEG analysis of AL_DCF versus ST, we identified 1139 DEGs, with 310 upregulated genes and 829 downregulated genes (**Figure 2. E)**. The upregulated genes were annotated with GO pathways related to developmental processes (GO:0032502), anatomical structure development (GO:0048856), cell differentiation (GO:0030154), cell adhesion (GO:0007155), cell migration (GO:0016477), chemotaxis (GO:0006935), and leukocyte chemotaxis (GO:0030595). The downregulated genes were annotated with various GO pathways related to tissue development (GO:0009888), epithelial development (GO:0060429), keratinization (GO:0031424), and cornification (GO:0070268) (**Figure 2. F)** (**Supplementary figure 4**).

Our results revealed a significant alteration in the expression of mechanosensing markers. Notably, we observed an augmentation in the expression of *PIEZO2*, while concurrently noting a reduction in the expression levels of *TRPV3*, *TRPV6*, and *OCLN* in both the AL and AL-DCF. Furthermore, a significant decrease was observed in the expression of lubricant mucins markers (*MUC16*, *MUC20*, *MUC4*, *MUC2*, and *MUC6*) in both AL and AL_DCF. In contrast, a notable increase was observed in the expression of genes related to inflammation, extracellular matrix (ECM) remodeling, and disruption (*IL6*, *CXCL2*, *CCL5*, *COL3A1*, *COL1A2*, *COL8A1*, *COL5A3*, *MMP1*, *MMP2*, and *MMP17*) in both AL and AL_DCF (**Figure 3. A, B)**.

**Figure 3.**
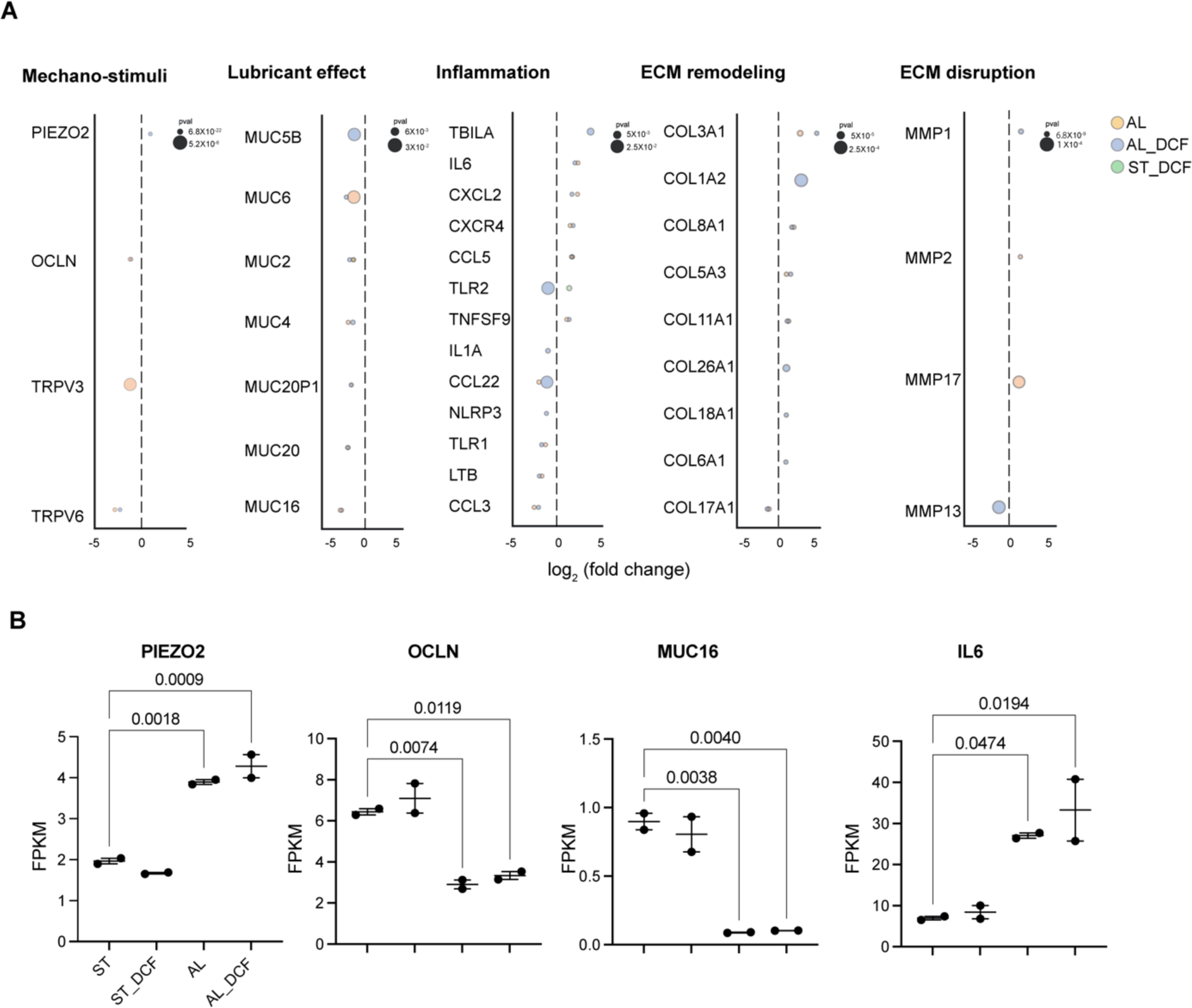
The impact of AL stimulus on the alteration key DE related transcripts. (A) DEGs of mechano-stimuli, lubricant effect, inflammation, ECM remodeling, and ECM disruption. Genes with 2-fold change and a *p*-value <0.05 was applied. (B) The comparative gene expression (normalized FPKM values) of DES markers in HCE-T cells. Dot plot in which the mean of each group is indicated with a black line (data are presented in duplicates as means ± S.E.M). The p-values were determined by the Dunnett’s multiple comparison test.

The introduction of DCF did not reverse of the overexpression of IL-6, nor did it mitigate the reduction in the gene expression of mucin and occudens, as exemplified by *MUC1* and *MUC16*, and *OCLN* (**Figure 3. B)**. Furthermore, the downregulation of the antioxidative enzyme related gene (*GSS*) remained unchanged despite the presence of DCF. However, it is noteworthy that the growth factor *TGFB1* transcript demonstrated a significant increase under AL stimuli. Intriguingly, this heightened expression was subsequently attenuated upon the addition of DCF in the AL-DCF treatment group (**Supplementary figure 5**).

### AL elevates mitochondria-related metabolites

To investigate the impact of AL and AL_DCF in inducing change in the extracellular metabolites in both the apical (AP) and the basolateral (BA) compartments, none-targeted-based LCMS metabolomic analysis was performed at 0, 3, 6, and 24 h (**Figure 4. A (I)**). We were able to annotate a total of 121 metabolites (**Supplementary data 1)**. We utilized peak areas for metabolite semi-quantification. The principal component analysis (PCA) results demonstrated a gradual shift in metabolite levels over time. This shift was evident from the systematic change in sample clusters at 3 h compared to 0 h, 6 h compared to 0 h, and 24 h compared to 0 h (**Figure 4. A (II)**). To identify unique metabolites, we employed partial least square discrimination analysis (PLS-DA) in combination with the variable importance in projection (VIP) (**Figure 4. B)**. We considered metabolites with a score equal to or greater than 2. At 3 h, the identified metabolites included methyl-2-oxovaleric acid, 3-methyl-2-oxobutanoic acid, palmitoyl-carnitine, myristoyl-carnitine, hippuric acid, ureidopropionic acid, cytosine, and N, N-dimethylguanosine. At 6 h, the identified metabolites were stearoyl-carnitine, palmitoyl-carnitine, methyl-2-oxovaleric acid, 3-methyl-2-oxobutanoic acid, myristoyl-carnitine, and ureidopropionic acid. Finally, at 24 h, the identified metabolites were 3-methyl-2-oxobutanoic acid, methyl-2-oxovaleric acid, lauroyl-carnitine, decanoyl-carnitine, thiamine, and 3-hydroxyisovaleric acid. In addition, we investigated the translocation of DCF from the AP compartment into the BA compartment. We calculated the accumulation of DCF in the BA compartment over a 24 h period based of the AUC values. Our results revealed a rapid accumulation of DCF in the BA compartment within the first 6 h. However, after this initial period, we observed a saturation point where the accumulation of DCF reached a plateau and remained relatively stable up to 24 h (**Supplementary figure 6**).

**Figure 4.**
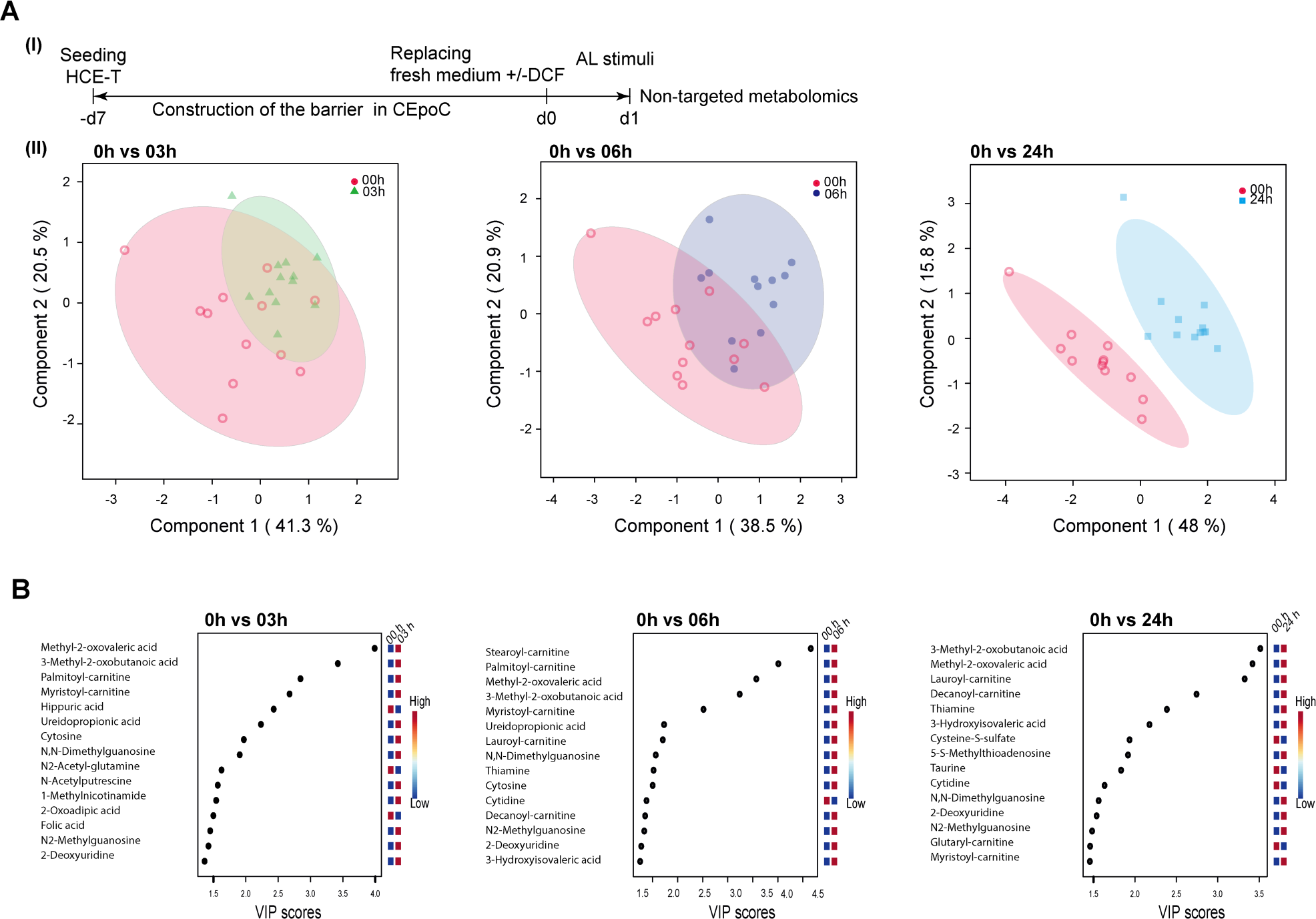
LCMS-based nontargeted metabolomic analysis of extracellular metabolites under AL stimulus. (A) I) The protocol that showcase the timeline of HCE-T culturing, barrier formation and extracellular samples collection (one microliter) for nontargeted metabolomics; II) Principal component analysis (PCA) of metabolomics dataset peak areas (log transformation) of samples (except internal standards, QC, and blanks) at 0 h vs 3 h, 0 h vs 6 h and 0 h vs 24 h, respectively. (B). Variable importance in projection (VIP) derived from the partial least square discrimination analysis (PLS−DA) that included all samples (except internal standards, QC, and blanks) for the selection of important metabolites at 0 h vs 3 h, 0 h vs 6 h and 0 h vs 24 h, respectively a p-value <0.05 was applied. Blue: downregulated genes, yellow: upregulated genes.

We subsequently directed our focus towards unique discovered metabolites, such as methyl-2-oxovaleric acid, 3-methyl-2-oxobutanoic acid, palmitoyl-carnitine, and decanoyl-carnitine. These metabolites were found to be secreted from both the AB and BA sides, albeit with distinct secretion kinetics.

To analyze the differences in secretion activity between AL and AL_DCF, we first adjusted the values based on channel volumes and then normalized them using the initial values at time 0. Next, we utilized the area under the curve (AUC) values to compare the metabolic activity. Under AL stimulation, both methyl-2-oxovaleric acid and 3-methyl-2-oxobutanoic were actively secreted from the AP side, exhibiting AUCs of 111.8 and 124.8, respectively. In comparison, their secretion activity on the BA side was lower, with AUCs of 44.79 and 39.91, respectively (**Figure 5. A (I)**). Interestingly, the combination of AL with DCF enhanced the secretion of these metabolites on the AP side, resulting in increased AUCs of 151.6 (methyl-2-oxovaleric acid) and 159 (3-methyl-2-oxobutanoic acid). The elevation in secretion was accompanied by a slight increase in BA secretion as well, with AUCs of 55.71 (methyl-2-oxovaleric acid) and 51.31 (3-methyl-2-oxobutanoic acid) (**Figure 5. A (II)**).

**Figure 5.**
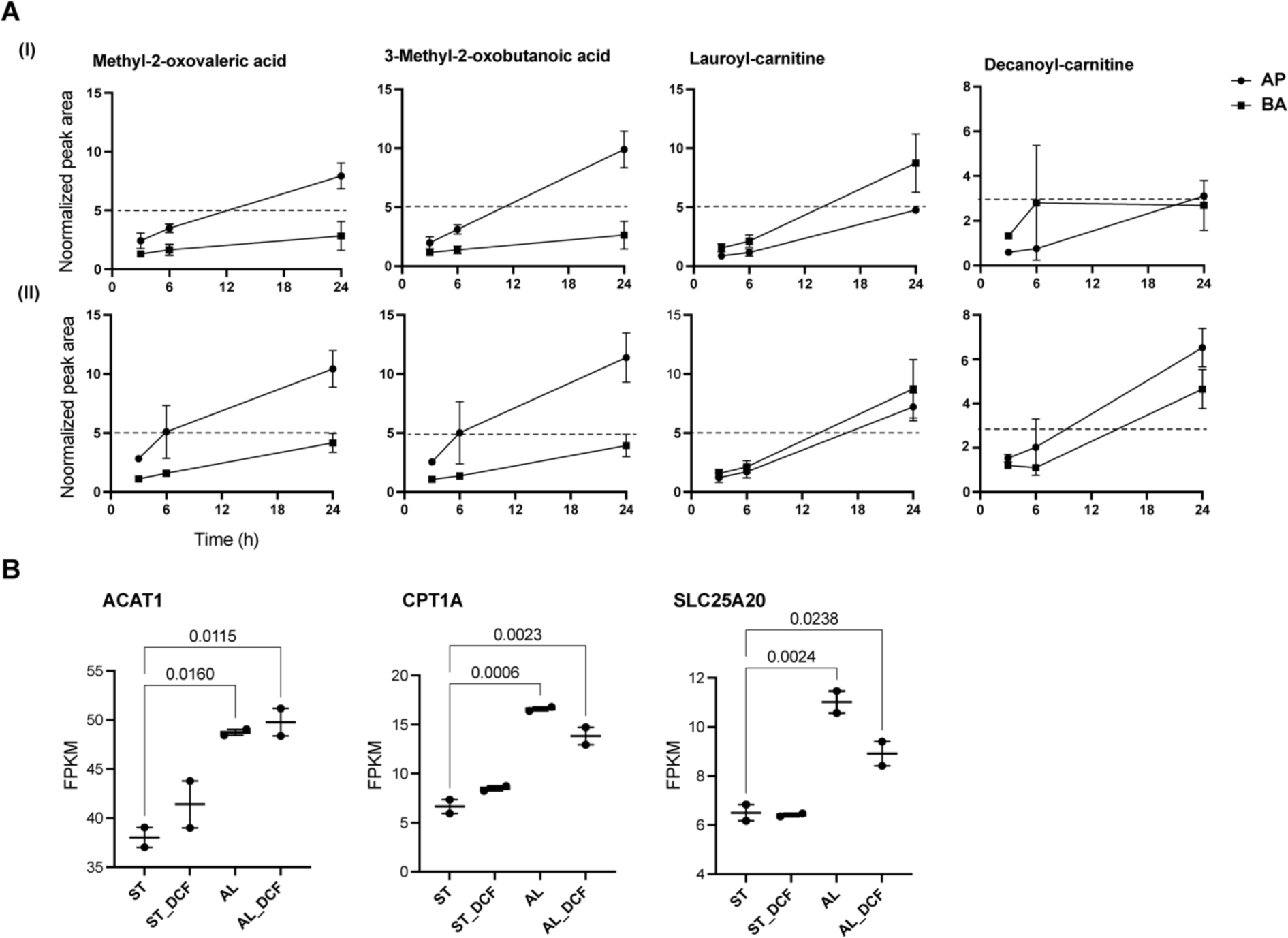
The effect of DCF on the alteration of extracellular metabolites under AL stimulus. (A) Selection of metabolites based on the VIP analysis score of highly active profile. I) under AL stimulus only; II) under AL stimulus and DCF treatment. Peak areas were corrected by the channels volume and then normalized by the 0 h values. AP: apical, BA: basolateral. Data are presented in triplicates as means ± S.E.M. (B) The comparative gene expression (normalized FPKM values) of essential metabolites-related genes in dot plot in which the mean of each group is indicated with a black line (data are presented in duplicates as means ± S.E.M). The *p*-values were determined using the Dunnett’s multiple comparison test.

On the contrary, in response to AL stimuli, lauroyl-carnitine and decanoyl-carnitine exhibited a tendency for active secretion on the BA side, showing AUCs of 103.4 and 55.63, respectively, in comparison to the AP side where their AUCs were 56.19 (lauroyl-carnitine) and 36.79 (decanoyl-carnitine). When combined with DCF, there was a slight increase in AP secretion, resulting in AUCs of 84.49 (lauroyl-carnitine) and 82.2 (decanoyl-carnitine). However, there was no notable secretion change in BA compartment, with AUCs of 103.4 and 55.26, respectively. We further investigated genes with biological connections to these metabolites. For example, the Acetyl-CoA Acetyltransferase 1 (*ACAT1*) gene encodes an enzyme that catalyzes the breakdown of acetoacetate, a product of both methyl-2-oxovaleric acid and 3-methyl-2-oxobutanoic acid. Interestingly, both AL and AL_DCF treatments led to a significant upregulation of this gene compared to the ST and ST_DCF treatments. In a similar manner, the Carnitine Palmitoyltransferase 1A (*CPT1A*) gene encodes an enzyme responsible for transferring long-chain fatty acids onto carnitine, forming acylcarnitines, as exemplified by lauroyl-carnitine and decanoyl-carnitine. Moreover, the Solute Carrier Family 25 Member 20 (*SLC25A20*) gene encodes a transporter accountable for conveying acylcarnitines across the mitochondrial membrane and governing mitochondrial oxidative metabolism. Our data reveals a noteworthy upregulation in the expression of *CPT1A* and *SLC25A20* genes under both AL and AL_DCF conditions as compared with ST and ST_DCF conditions. Notably, the AL_DCF group demonstrated a slightly diminished expression of *SLC25A20* (**Figure 5. B**).

## Discussion

DES originates from various factors, such as the instability of the tear film and inadequate tear secretion. These factors expose the ocular surface to air, potentially resulting in tissue damage. To replicate the conditions of evaporative dry eye, we recognized the significance of applying an air-liquid interface stimulus (AL) directly to the surface of the corneal epithelial barrier for inducing dry eye-like characteristics in cells.

In our study, we thought to adapt our previously developed CEpoC by implementing additional modifications for this specific purpose. As part of these modifications, we introduced additional reservoirs in the outlet areas. This enhancement was aimed at increasing the exposure of cells to air. Furthermore, we incorporated a bi-directional flow stimulus, while periodically introducing air into the cell chamber at regular intervals.

Subsequently, we proceeded to examine the formation of the corneal epithelial barrier within the adapted CEpoC using the method we had previously employed with HCE-T cells. Our findings revealed that over the course of a week, HCE-T cells grew and expanded within the device, ultimately leading to the development of a barrier. The functionality of the barrier was further verified through the observation of tight junction protein (ZO-1) and other key markers such as P-gp and CK12. Collectively, these findings underscored the feasibility of creating a human corneal epithelial barrier within the adapted device.

We proceeded to apply the AL stimulus to the CEpoC for a duration of 24 h. We then assessed cell viability and morphology under two conditions: AL alone, and AL in combination with DCF (50 μM), as compared to cells under static (ST) conditions. We decided the concentration of DCF of 50 μM as previously reported^13^, while a higher dose (200 μM) resulted in considerable cell death (data is not presented). Our results demonstrated that the exposure to the AL stimulus did not lead to notable changes in cell viability, a conclusion supported by the Calcein AM intensity measurements. Moreover, the administration of DCF treatment did not trigger any further alterations. It is worth noting that the use of DCF in the treatment of DES has generated conflicting outcomes in the literature. While it has been proposed as a potential treatment for DES ^13^, there have been contrasting reports highlighting cases of toxic reactions in corneal epithelial cells following DCF application^14,15^. In our results, we did not observe significant toxicity as determined by the Calcein AM assay. To deepening our understanding about that, we then investigated the transcriptomics alternation under AL and AL_DCF.

We observed a distinct transcriptomic change between cells exposed to AL and AL_DCF conditions compared to those under ST and ST_DCF conditions, signifying a fundamental shift in the transcriptomic expression within the cells. Our analysis of GO pathways revealed significant impacts on processes related to development, anatomical structure formation, DNA packaging, epithelium development, cornification, and cell-cell junctions in both AL and AL_DCF conditions. Furthermore, the pathway related to chemotaxis and leukocyte chemotaxis was affected specifically in the AL_DCF group. Remarkably, these pathways collectively mirror common characteristics of biological pathways associated with DES, where disruptions to epithelial structure, compromised cell-cell junctions, the initiation of inflammatory responses, and the remolding of ECM are prevalent features^16^. Our data revealed an increase in genes associated with inflammation (such as *IL6*) and a substantial decrease in genes linked to mucin production, notably *MUC16*. These results align with common observations in cases of DES. Furthermore, the AL stimulus elicited a noteworthy upregulation of *PIEZO2*, a mechanosensing channel expressed in the corneal epithelium^17^, which could exacerbate the signaling of damage in DE patients^18^. Conversely, we observed a downregulation in the expression of *OCLN*, a gene related to cell-cell junctions. This indicates that the AL stimulus could indeed disrupt tight junctions as anticipated in DES conditions^19^. Although TRP channels, such as *TRPV3*, are expressed in the cornea and are associated with thermosensation^20^, their contribution to DES is likely minor. As a result, we did not observe a significant increase in their expression in our data.

Non-targeted metabolomic analysis of extracellular metabolites revealed a significant secretion of acylcarnitines, particularly on the BA side, in response to AL and AL_DCF, which aligns with our previous findings^11^. Intriguingly, the addition of AL_DCF led to an unexpected kinetic shift, increasing the secretion of these metabolites in the AP direction. It is worth noting that excessive secretion of acylcarnitines, such as lauroyl-carnitine, has been associated with inflammation ^21^. Furthermore, both methyl-2-oxovaleric acid and 3-methyl-2-oxobutanoic were prominently secreted apically, with higher levels observed in the AL_DCF condition. These metabolites are considered metabotoxins, indicating an underlying disruption in cellular metabolism ^22^.

Collectively, our results suggest that AL stimulus enhances the secretion of metabolites indicative of cell damage and inflammation. This observation correlates with the upregulation of genes associated with inflammation (e.g., *IL-6*) and cellular damage (e.g., *OCLN* and *MUC16*). Interestingly, the addition of DCF does not appear to effectively mitigate these effects, potentially explaining the undefined outcomes observed in DES thus far. Notably, there is a slight downregulation of *TGFB1* gene expression under the combined influence of DCF and AL. TGFB1 secretion is known to play a crucial role in cornea injury responses and the activation of keratinocytes in the corneal stroma ^23^. This suggests that DCF may modulate corneal wound healing signaling rather than interfering with inflammation pathway.

In conclusion, we examined how stimulating the air-liquid (AL) interface affects corneal epithelial cells in CEpOC with a purpose of mimicking conditions seen in evaporative dry eye. We successfully upregulated the CEpOC device to create an AL interface. By using HCE-T cells, we cultivated a functional corneal epithelial barrier within the modified CEpOC model over a period of seven days. Transcriptomic analysis unveiled distinct patterns of gene expression in response to AL and AL_DCF stimuli. These changes mirrored characteristics of DES, such as disrupted epithelial structure and inflammation. Metabolomic analysis revealed the secretion of metabolites, particularly acylcarnitines, in response to AL and AL_DCF stimuli. Interestingly, DCF increased the apical secretion of these metabolites, which are associated with cellular damage and inflammation. Our study sheds light on the molecular and metabolic changes that underlie AL stimuli. The observed alterations suggest a cascade of events involving inflammation, disrupted cellular structure, and altered metabolism. Importantly, our findings raise questions about the effectiveness of DCF in mitigating these effects. Future studies can further examine the specific signaling pathways and mechanisms driving the observed changes. Additionally, exploring the long-term effects of AL stimulus and DCF treatment on ocular health and therapy is needed.

## Methods

### Microfluidic device fabrication

The microfluidic device was fabricated using stereolithographic 3D-printing techniques and solution cast-molding processes. Briefly, a mold for the microfluidic channels was produced using a 3D printer (Keyence Corporation, Osaka, Japan). Two molds were fabricated: upper blocks and lower blocks. Each block contained four chambers (15 mm length, 1.5 mm width, and 0.5 mm height). Prior to use, molds surfaces were coated with trichloro(1H,1H,2H,2H-perfluorooctyl) silane (Sigma-Aldrich, St. Louis, MO, USA). A Sylgard 184 PDMS two-part elastomer (ratio of 10:1 pre-polymer to curing agent; Dow Corning Corporation, Midland, MI, USA) was mixed, poured into the molds to produce a 4-mm-thick PDMS upper layer and a 0.5-mm-thick PDMS lower layer, and degassed using a vacuum desiccator for 1 h. The PDMS in the lower block was fixed on a glass slide. The PDMS material was then cured in an oven at 80°C for 24 h. After curing, the PDMS was removed from the molds, trimmed, and cleaned. A clear polyester (PET) membrane (pore size of 0.4 μm, thickness 10 μm, nominal pore density of 4 × 10^6^ pores cm ^2^) was fixed on each chamber of the lower PDMS block. Both PDMS blocks were corona-plasma-treated (Shinko Denki, Inc., Osaka, Japan) and bonded together by baking in an oven at 80°C.

### Human corneal epithelial cell culture

The human corneal epithelial cell line (HCE-T) was obtained from the RIKEN Bioresource Research Centre (Ibaraki, Japan). HCE-T cells were grown in a culture medium of DMEM/F12 supplemented with 5% fetal bovine serum, 5 μg/mL insulin, 10 ng/mL human epithelial growth factor, and 0.5% dimethyl sulfoxide. The cells were passaged at a ratio of 1:4 using trypsin-EDTA solution.

### The construction of corneal epithelial barrier in the microfluidic device

Before use, the microfluidic cell culture device was sterilized by placing it under ultraviolet light in a biosafety cabinet for 30 minutes. The channels of the device were then washed with the DMEM/F12 medium. Cells were harvested using trypsin and resuspended in DMEM/F12. The cell suspension was introduced into the upper channel of the device through a cell inlet with a cross-sectional area of 0.23 cm^2^, at a density of 1x10^6^ cells per mL. The microfluidic device was placed in a humidified incubator at 37°C and5% CO^2^ for 7 days, with the medium in each chamber was changed every 24 h.

### The application of flow stimulus

Prior to the initiation of the AL stimulus, microfluidic channels were filled with fresh culture medium. For drug treatment with DCF, DCF was added to the upper channels at a final concentration of 50 μM, with a volume of 15 μL in the channel and 20 μL in the reservoir. The upper channels were exposed to the AL flow using a syringe pump (KD Scientific, Fisher scientific, Holliston, USA). Sterilized tubes (Tygon LMT-55) were divided into eight branches, which were then connected to an air reservoir tube. The entire setup was linked to a syringe pump. The dynamic flow model consisted of three phases: Phase 1: withdrawal flow at a rate of 100 μL/s. Phase 2: infusion flow at a rate of 100 μL/s. Phase 3: a pause. Throughout these phases, lower channels were maintained under static conditions.

### The determination of cell viability

Cell viability was assessed by the live staining with Calcein AM (Dojindo Molecular Technologies, Inc.). Briefly, cells were incubated with Calcein AM at a final concentration of 10 μg mL^-1^ in DMEM/F12 medium at 37°C for 60 min. The cells were then washed twice with PBS- and subjected to microscopic imaging.

### Immunofluorescence and microscopic imaging

For the immunostaining, cells were fixed with 4% paraformaldehyde in PBS-for 25 min at 25°C and then permeabilized with 0.5% Triton X-100 in PBS-for 10 min at 25 °C. Subsequently, cells were blocked with blocking buffer (5% (v/v) bovine serum albumin, 0.1% (v/v) Tween-20) at 25 °C for 1 h and then incubated at 4 °C for overnight with the primary antibody in blocking buffer (**Supplementary table 1)**. Cells were then incubated at 37°C for 60 min with a secondary antibody (Alexa Fluor 488 donkey anti-rabbit IgG and Alexa Fluor 594 donkey anti-mouse IgG 1:1000; Jackson ImmunoResearch, West Grove, PA, USA) in blocking buffer prior to a final incubation with 4′,6-diamidino-2-phenylindole (DAPI) or anti-phalloidin for F-actin (Invitrogen) at 25 °C. For imaging, we used a Nikon ECLIPSE Ti inverted fluorescence microscope equipped with a CFI plan fluor 10×/0.30 N.A. objective lens (Nikon, Tokyo, Japan). Images were then analyzed using ImageJ software (National Institute of Health, Maryland, USA). Cell Profiler software (Version 3.1.8; Broad Institute of Harvard and MIT, USA^19^).

### RNA extraction and sequencing

RNA extracted using RNeasy Mini Kit (Qiagen, Hilden, Germany) according to the maker instruction. Samples quality was assured by using the bioanalyzer (Agilent technologies, Inc., USA) with an RNA Integrity Number (RIN) value greater than or equal or higher than 7. Samples were then applied for the next generation sequencing (Macrogen, Tokyo, Japan) starting with RNA-seq library construction (TruSeq stranded mRNA LT Sample Prep Kit) followed by sequencing by NovaSeq 6000 Illumina system using 101 reads per specimen.

### RNA-seq data mining and GO enrichment analysis

The quality control of the sequenced raw reads was performed by calculating overall reads’ quality, total bases, total reads, GC (%) and basic statistics (FastQC v0.11.7). In order to reduce biases in analysis, artifacts such as low-quality reads, adaptor sequence, contaminant DNA, or PCR duplicates were removed (Trimmomatic0.38)^24^. Trimmed reads were mapped to reference genome with HISAT2, splice-aware aligner (HISAT2 version 2.1.0, Bowtie2 2.3.5.1)^25,26^. Transcript was assembled by StringTie with aligned reads (StringTie version 2.1.3b)^27^. This process provided information of known transcripts, novel transcripts, and alternative splicing transcripts. Expression profiles were represented as read count and normalization value which is based on transcript length and depth of coverage. The FPKM (Fragments Per Kilobase of transcript per Million Mapped reads) value or the RPKM (Reads Per Kilobase of transcript per Million mapped reads) is used as a normalization value. Differentially Expressed Genes (DEG) analysis was conducted using read count values obtained from StringTie. Initially, low-quality transcripts were filtered out during data preprocessing. Then, (Trimmed Mean of M-values) TMM normalization was applied. For statistical analysis, Fold Change and exact Test from edgeR were employed for each comparison pair. Significant results were identified based on the criteria of |Fold Change| >= 2 and an exact Test raw p-value < 0.05. Enrichment test was conducted with significant gene list using g:Profiler tool platform ^28^.

### Extracellular metabolites collection and analysis by LCMS

One microliter of extracellular samples was collected as previously reported^112930^. Tubes containing one microliter of dried extracellular samples were thawed and 150 μL of water and acetonitrile (3:7, v/v) was added, containing three technical internal standards (tISs): 0.1 µM CHES, 0.1 µM HEPES, and 0.2 µM PIPES. After resuspension, all samples were centrifuged at room temperature for 1 min at 1000× g. A quality control (QC) sample was prepared by pooling an aliquot from each sample. Next, 40 μL of the supernatant was transferred to a 96-well 0.2 mL PCR plate (PCR-96-MJ; BMBio, Tokyo, Japan). The plate was sealed with a pierceable seal (4titude; Wotton, UK) for 3 s at 180 °C using a plate sealer (PX-1; Bio-Rad, Hercules, CA, USA) and maintained at 10 °C during the LC-MS measurements. The LC-MS method has been described previously^31^. The injection volumes were 7 μL in negative and 3 μL in positive ionization mode. In brief, metabolite separation was achieved on an Agilent 1290 Infinity II system using a SeQuant ZIC-HILIC (Merck, Darmstadt, Germany) column and a gradient between water containing 0.1% formic acid (pH = 2.6) and acetonitrile containing 0.1% formic acid in positive ionization mode, and a SeQuant ZIC-pHILIC (Merck, Darmstadt, Germany) column and a gradient of acetonitrile and 5 mM ammonium acetate in water (pH = 9.3) in negative ionization mode. Data were acquired on an Agilent 6550 Q-TOF-MS system with a mass range of 40−1200 m/z in all ion fragmentation mode, including three sequential experiments at alternating collision energies: full scan at 0 eV, followed by MS/MS scans at 10 eV and 30 eV, with a data acquisition rate of 6 scans/s. Data were converted to mzML format using Proteowizard and processed using MS-DIAL version 4.80^32,33,34^ An in-house library containing accurate masses (AMs) and retention times (RTs) for 622 compounds obtained from chemical standards was used to annotate the detected compounds. Peak areas exported from MS-DIAL were used for metabolites’ semi-quantification. Only metabolites with a coefficient of variation (CV) less than 30% in the QC samples or D-Ratio^35^ <50 were used for further analysis.

### Data visualization and statistics

Data are represented as mean ± S.E.M. RNA-seq data are presented in duplicate of biological samples. Other data are presented of triplicate or more of biological samples. The unpaired t test, Tukey’s, Dunnett multiple comparison test were performed using GraphPad prism 8 (GraphPad Software, La Jolla California, USA). Principal component analysis (PCA) visualization was performed using the MetaboAnalyst platform^36^. Orange 3 software (Version 3.23.1; Bioinformatics Laboratory, Faculty of Computer and Information Science, University of Ljubljana, Slovenia^12^) were used for data mining. Python Jupyter notebook 6.1.4, with Pandas and Bioinfokit packages. Data visualization was performed using Python 3 with matplotlib and seaborn packages.

## Data availability

RNA-seq data have been deposited to the GEO repository under GEO accession number GSE244787.

## Acknowledgments

Funding was generously provided by the Japan Society for the Promotion of Science (JSPS, 22K14548 and JSPS, 20KK0160), and the Hirose Foundation to Rodi Kado Abdalkader. We acknowledge the WPI-iCeMS supported by the World Premier International Research Centre Initiative (WPI), the Ritsumeikan Global Innovation Research Organization (R-GIRO) and Gunma University Initiative for Advanced Research (GIAR) for their support.

## Authors contribution statement

Rodi Kado Abdalkader took responsibility for the overarching research concept, project management, biological experiment design and execution, data analysis and interpretation, results visualization, and manuscript writing. Romanas Chaleckis conducted the non-targeted metabolomic LC-MS analysis, contributing to data interpretation and manuscript refinement. Ken-ichiro Kamei and Takuya Fujita provided resources and contributed to data interpretation and manuscript improvement. All authors have reviewed and approved the final version of the manuscript for publication.

## Additional information

Competing interests: All authors declare no competing financial interest.

## Supplementary Information

**Supplementary Figure 1.**
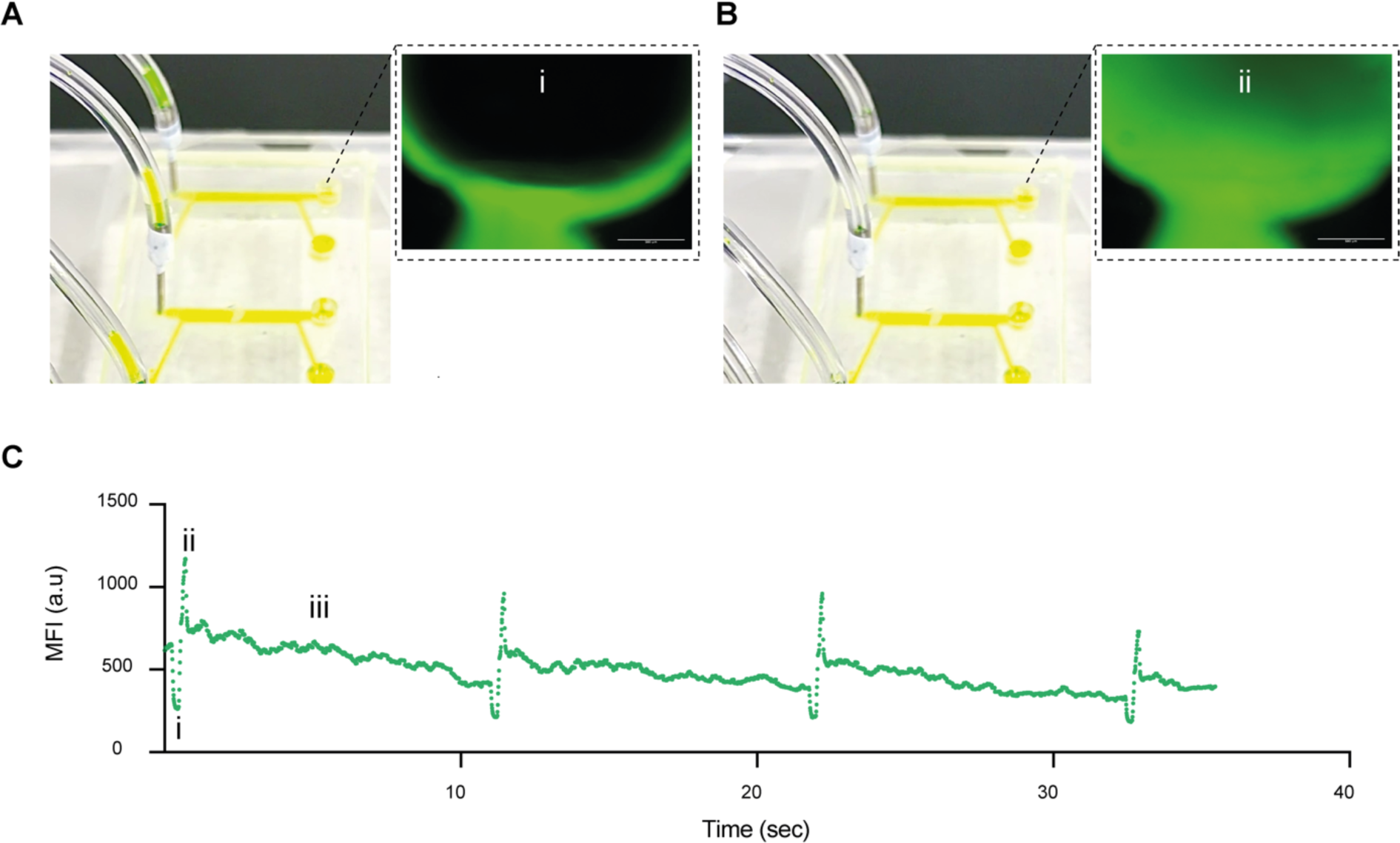
The process of AL stimulus under cell-free conditions. (A) Air withdrawal (i) (A) Air infusion (ii). The green signal is captured from the sodium fluorescein solution. (C) Graph that showcase the mean fluorescence signal in the reservoir area during air withdrawal phase (i), air infusion phase (ii), and pause phase (iii).

**Supplementary Figure 2.**
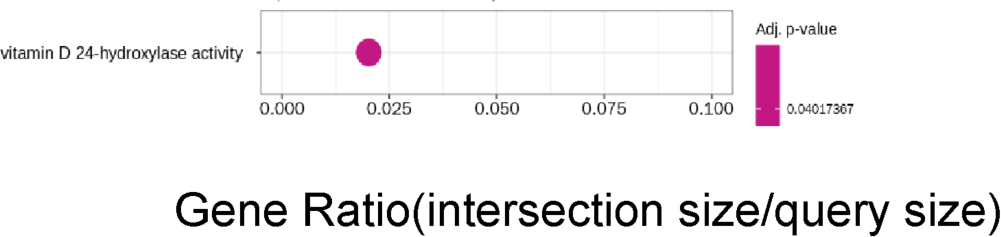
GO pathways of DEGs in of ST_DCF vs ST. Genes with 2-fold change and a p-value <0.05 was applied.

**Supplementary Figure 3.**
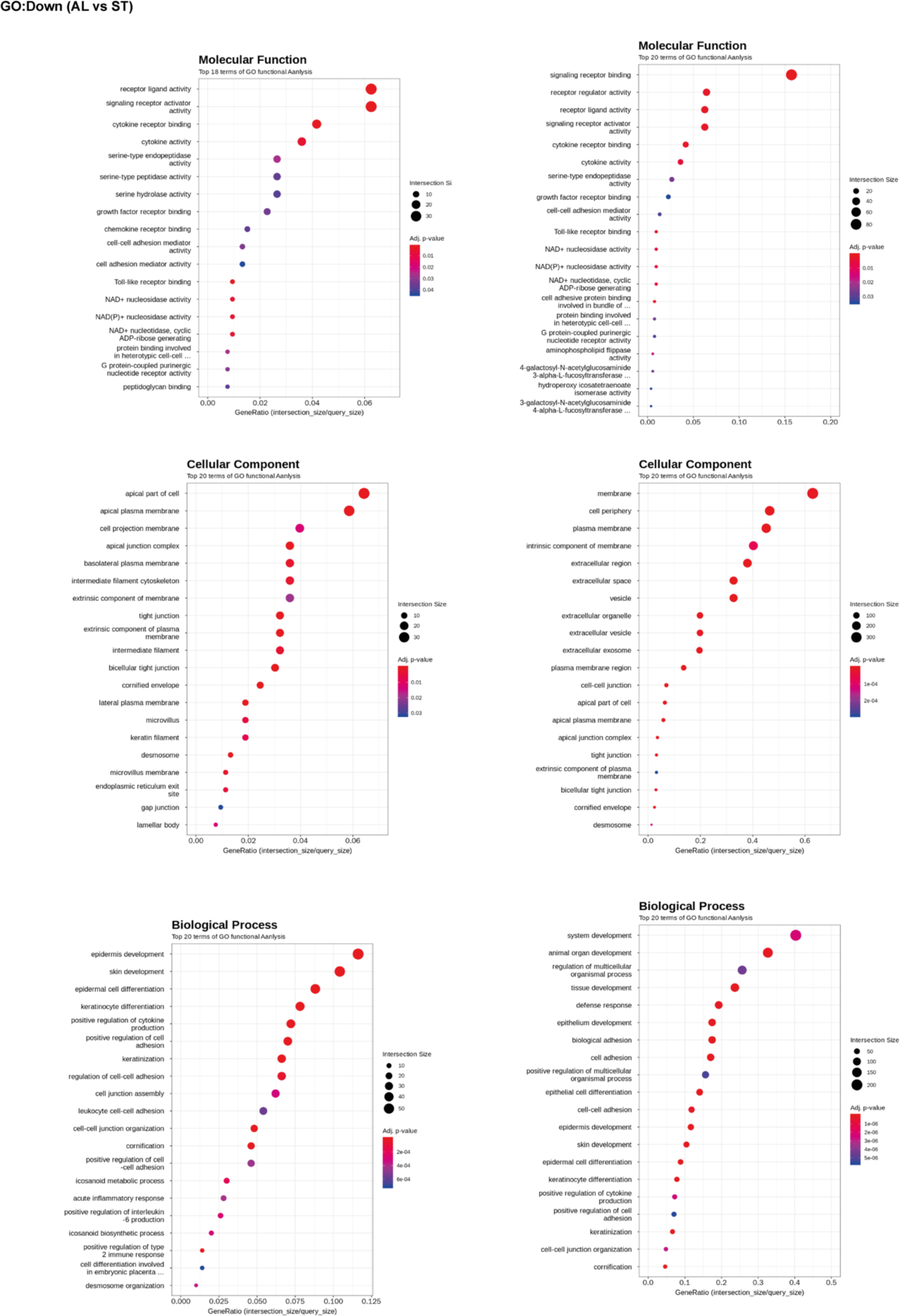
GO pathways of DEGs in of AL vs ST. Genes with 2-fold change and a p-value <0.05 was applied.

**Supplementary Figure 4.**
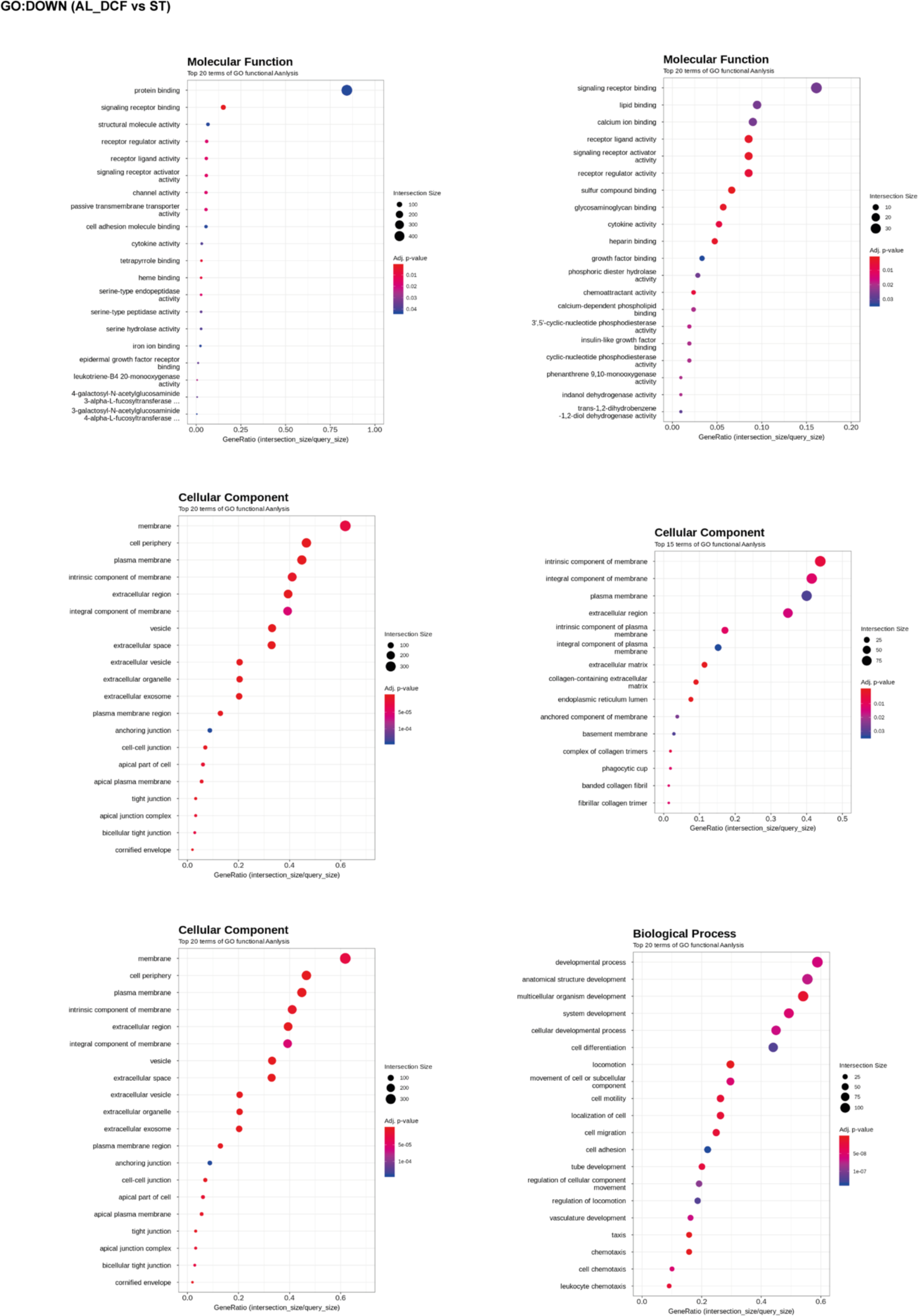
GO pathways of DEGs in of AL_DCF vs ST. Genes with 2-fold change and a p-value <0.05 was applied.

**Supplementary Figure 5.**
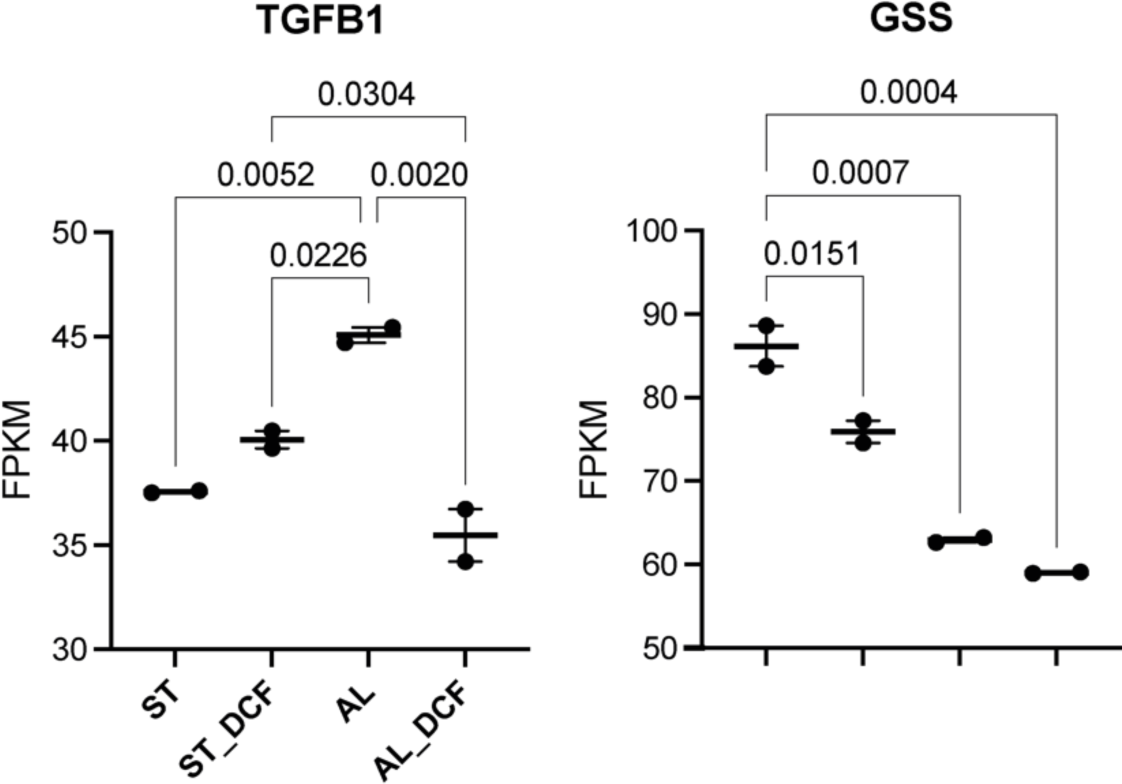
The comparative gene expression (normalized FPKM values) in dot plot in which the mean of each group is indicated with a black line (data are presented in duplicates as means ± S.E.M). The *p*-values were determined using the Tukey HSD test and Dunnett’s multiple comparison test.

**Supplementary Figure 6.**
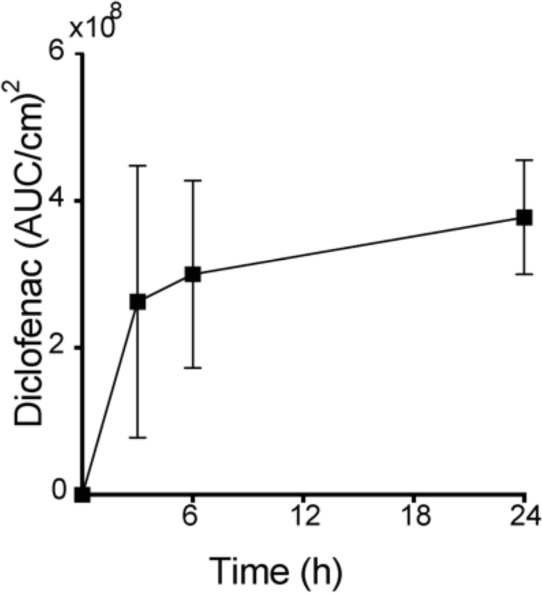
Diclofenac (DCF) accumulation in the basolateral compartment. **Data are presented in triplicates as means ± S.E.M.**

**Supplementary Table 1.** List of reagents and resources.

| Item | Maker | Catalog number | Details |
| --- | --- | --- | --- |
| ZO-1 antibody, Rabbit | Thermo Fisher | 61-7300 | IF 1:50 |
| P-gp antibody, Rabbit | Abcam | ab129450 | IF 1:50 |
| CK12 antibody, Mouse | Santacruz | sc-515882 | IF 1:200 |
| Diclofenac sodium | Wako | 043-22851 |  |

